# Mechanical influences on *in silico* tumor evolution

**DOI:** 10.1101/2021.03.23.436580

**Authors:** Jakob Rosenbauer, Marco Berghoff, James A. Glazier, Alexander Schug

## Abstract

Experimental insight and conceptual understanding of tumor growth are steadily growing and leading to new therapeutic interventions. Experiments and clinical studies are able to link single-cell properties to macroscopic tumor attributes. The development of cellular subpopulations in heterogeneous tumors can be understood as an evolutionary system with different cell types competing over both space and nutrients. However, to predict the growth trajectory and development of a tumor, fitness and trade-offs of cell properties in the context of the surroundings are required and often inaccessible. The optimum of the evolutionary trajectory provides a target for intervention, but can mostly not be identified. We propose that the optimal value of cellular properties is influenced by the tumor surrounding. Computational multiscale-modeling of tissue enables the observation of the trajectory of each cell while modeling the tumor surrounding. We model a 3D spheroid tumor and the fitness of individual cells and the evolutionary behavior of the tumor are quantified and linked to global parameters. Cell–cell adhesion and cell motility are two important mechanical properties for cell development and used as free parameters. Mechanical properties alone are able to drive the tumor towards low adhesion.We implement a dynamically changing nutrient surrounding representing the fluctuating blood-supply through blood vessel collapse and angiogenesis. We find that the evolutionary speed depends on the frequency of the fluctuations. We identify a frequency domain in which the evolutionary speed is significantly increased over a tumor with constant nutrient supply. The findings suggest that mechanically-induced fluctuations can accelerate tumor evolution.

**Author summary:** Limited space and nutrients together with competing cell types drive an evolutionary process inside tumors. This process selects for the fittest cell types and optimizes the growing behavior for its local surroundings. An expanding tumor exerts mechanical forces on its cells and its surroundings, leading to a fluctuating nutrient supply through collapsing blood vessels. Here, we observe the influence of a dynamically changing surrounding on the evolutionary behavior of heterogeneous tumors in a high-resolution computational model. We find that the evolutionary speed depends on the frequency of the fluctuations and a fitness advantage of low-adhesion cells.

## Introduction

The emergence and development of tumors in humans still presents a significant challenge to medicine and is one of the leading causes of death. Knowledge about the development of tumors is continuously expanding. Especially, the large variety of tumor types, differences between patients and tumors consisting of many types of tumor cells (termed tumor heterogeneity) present challenges. Here, a tumor in surrounding solid tissue and the development of tumor properties is observed over time in a computational model. The development of a tumor can be described as an evolutionary system in which cell types present species competing over resources [1, 2]. The evolution of a tumor is driven by mutations. These mutations change the properties (e.g., motility or cell-cell adhesion) of cells, which then can provide advantages or disadvantages for further growth. Competition over resources and space leads to the selection of the most advantageous tumor cell types, whose subpopulation will overtake the rest. However, cells can not optimize all parameters at the same time, several trade-offs in cellular properties in cancer have been described and studied [3, 4]. We assume a limited energy budget for each cell and introduce a trade-off of proliferation against motility for cell types.

While the properties and parameters of a cell are accessible in experiments, the fitness of a cell is very hard to determine experimentally. The fitness determines the ability of a cell to reproduce in its surroundings and multiply, therefore cells with high fitness are selected for in evolution. To determine the fitness of cells experimentally, the lineage of cells has to be tracked over time, making it necessary to track each individual cell over a long time, which is currently not possible in *in vivo* tumors. Models of the fitness landscape based on genetic changes and driver mutations have been introduced [5], yet phenotypic changes in cells can also drive mutation [6, 7]. Naturally, an ensemble of cells with different cell types evolves towards the cell type of highest fitness. The fitness is not an intrinsic property of the cell but results from the interplay of cells with their surroundings. Finding the fitness of cell types depending on their parameters and surroundings allows for directed interventions to change the course of evolution. Taking control of the evolution of a tumor could serve as a tool to stop the spreading and further growth of a tumor and aid treatment.

The mechanical interactions of the tumor with its environment and between the tumor cells influence the trajectory of the tumor [8]. Cells with low cell–cell adhesion mechanically sort to the outside of tumors, providing a higher nutrient supply and therefore evolutionary advantages. The nutrition levels on the surface of tumors are higher than in the center of the tumor, leading to an induced advantage of low adhesion cells. Therefore, we will test this prediction and observe whether those cells have an evolutionary advantage in an evolving heterogeneous tumor. Cell motility describes the active movement of cells, we predict that random cellular movements should provide no significant evolutionary advantage in a constant surrounding. Phenotypical changes towards higher motility cells and the epithelial to mesenchymal transition are associated with tumor invasion [9–11]. In a dynamically changing nutrient environment, cells with higher motility can dynamically occupy the most advantageous positions. Enhanced motility is known to be a driver of the formation of metastases [12], therefore a dynamic nutrient field could induce metastases.

In this work, we focus on two variables in cellular properties, namely *cell–cell adhesion* and *motility*. Increased motility is a landmark for the development of metastases [12]. We hypothesize that mechanical and geometric constraints alone are sufficient to drive the evolution of a tumor towards high motility.

The computational modeling of tumor development and heterogeneity has evolved and was successfully applied, e.g., a cellular automaton model was used to observe the influence of cell dispersal and turnover on tumor heterogeneity [13]. The use of computational models in clinical applications and personalized treatment plans is gaining momentum [14, 15]. Work on tumor internal evolution has been recently done by Büscher et al. [16] where they compared adhesion and ‘growth strength’ in an evolving tumor and found that in some cases a mixture of different cell types emerges as a stable state. The importance of cell–cell adhesion for tumor invasion has been recently highlighted by Ilina et al. [17]. In Ref. [18] the authors showed with a mathematical model using the evolutionary game theory that tumor heterogeneity and the optimal cell phenotype depends on the microenvironment and position within the tumor. The ‘go-or-grow’ hypothesis describes reversible phenotype changes between motile and proliferative phenotypes in cancer. This hypothesis was proposed and tested with positive [19] and negative correlations [20] for different cancer cell lines. Trade-offs between two or more cell properties have been studied by Gallaher et al. [4]. Here, we are not focusing explicitly on the ‘go-or-grow’ hypothesis, since we assume each cell to retain its phenotype throughout its lifetime and changes of cell properties can only occur at mutation events during cell division.

Cell adhesion and cellular motility have been recently shown to strongly influence the invasion behavior of breast cancer and to differentiate between solid-like, fluid-like, and gas-like behavior [17]. Confinement by the extracellular matrix and cellular motility were recently investigated systematically and a phase space of tumor invasion modes was proposed [21].

The intricate surrounding around a developing tumor is important and strongly influences its progression together with the intrinsic properties of the tumor cells. An invading tumor is expanding in volume and requires nutrients that can only diffuse through a finite length of tissue. Angiogenesis, the growth of new blood vessels, is initiated for a better nutrient supply. This leads to a quick volume increase of the tumor, which builds up pressure and solid stresses within the tumor that can collapse blood vessels [22]. This interplay of formation and collapse of blood vessels can lead to a fluctuating availability of nutrients for the cells in the tumor.

During the development of tumors, single cell effects are of major importance, with mutations initially occuring in a single cell. Stochasitcity and rare events are non-negligible during tissue development. As we showed in [23], single cell effects are important for the patterning of tissue. Cell migration is facilitated by many concurrent processes and represents a multiscale process, therefore requiring multiscale modeling [24]. To capture the complexity of tumor growth, computational models with single cell resolution enable the incorporation of single cell behavior that would be difficult to capture using continuous ODE models. Since evolution is a stochastic process requiring many iterations to find a stochastically meaningful result and the requirement of large-scale tissues makes the use of high-performance computing necessary.

The recently developed framework CellsInSilico [25] is used for the simulations, it is based on the cellular Potts model (cpm) [26] that has been established for the simulation of tumor growth [27]. Through its parallelization and optimization for supercomputers, the framework enables large-scale three-dimensional simulations and high numbers of simulations, which is necessary to generate sufficient statistics to study the proposed problems.

We implement a discrete evolution of two independent parameters, namely cell-cell adhesion and cell motility. Cells can alter and change their parameters stepwise, with a small probability at each division. Parameters are changed incrementally, which implies a continuous evolution and neglects possible mutations with larger effects. For cell motility, we introduce a trade-off on division rates since we assume cells have a limited energy contingent. We simulate a spheroid tumor in an external gradient of nutrients. The tumor is surrounded by a population of non-mutating host tissue. We observe the evolution of the tumor composition along both free parameters. Different mechanisms that couple cell division rates to nutrient availability are compared. We compare the effect of temporally changing nutrient availability.

## Results

### Constant environment

First, we investigate the dynamics of the system without the dependency of cell division and death on nutrient availability. Division and death are determined by the respective rates and age and volume constraints. A spheroid tumor develops from an initial tumor seed, consisting of ≈ 3700 tumor cells (cf. Fig. 1) in a surrounding of nondividing and nondying cells. The tumor grows until the space provided by the 3D simulation box is used up by the surrounding tissue and the tumor. Cell division is limited to cells above a threshold volume (*V*_THRS_ = 0.9 *V*_0_) and cells are only compressible to a finite extent, therefore the absolute number of tumor cells is limited and the cells compete over the available space. In the emerging steady state, the tumor size remains constant with cells dying and dividing at equal rates. The cell division rate is higher than the death rate (cf. Fig. 2 a-c) and the absolute number of cells is geometrically constrained, thereby the effective division rate is limited by the death rate.

**Fig 1.**
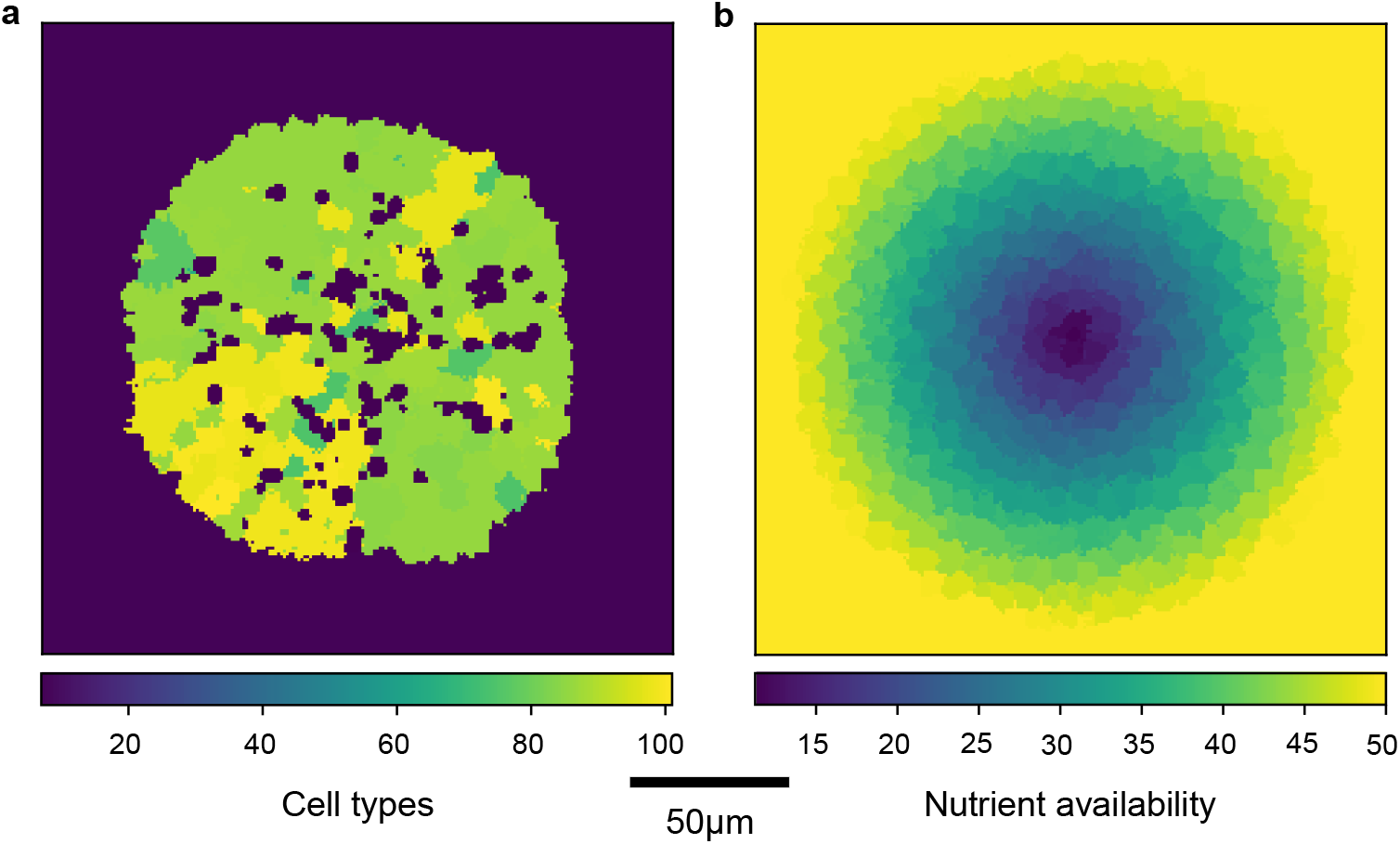
Geometry of tumor simulations. **a** Spatial organization of the cell types, 2D slice through the center of the 3D simulation. The color map indicates the different cell types, that are linearly numbered. **b** coloring by nutrient availability. Nutrient availability in arbitrary units (AI).

**Fig 2.**
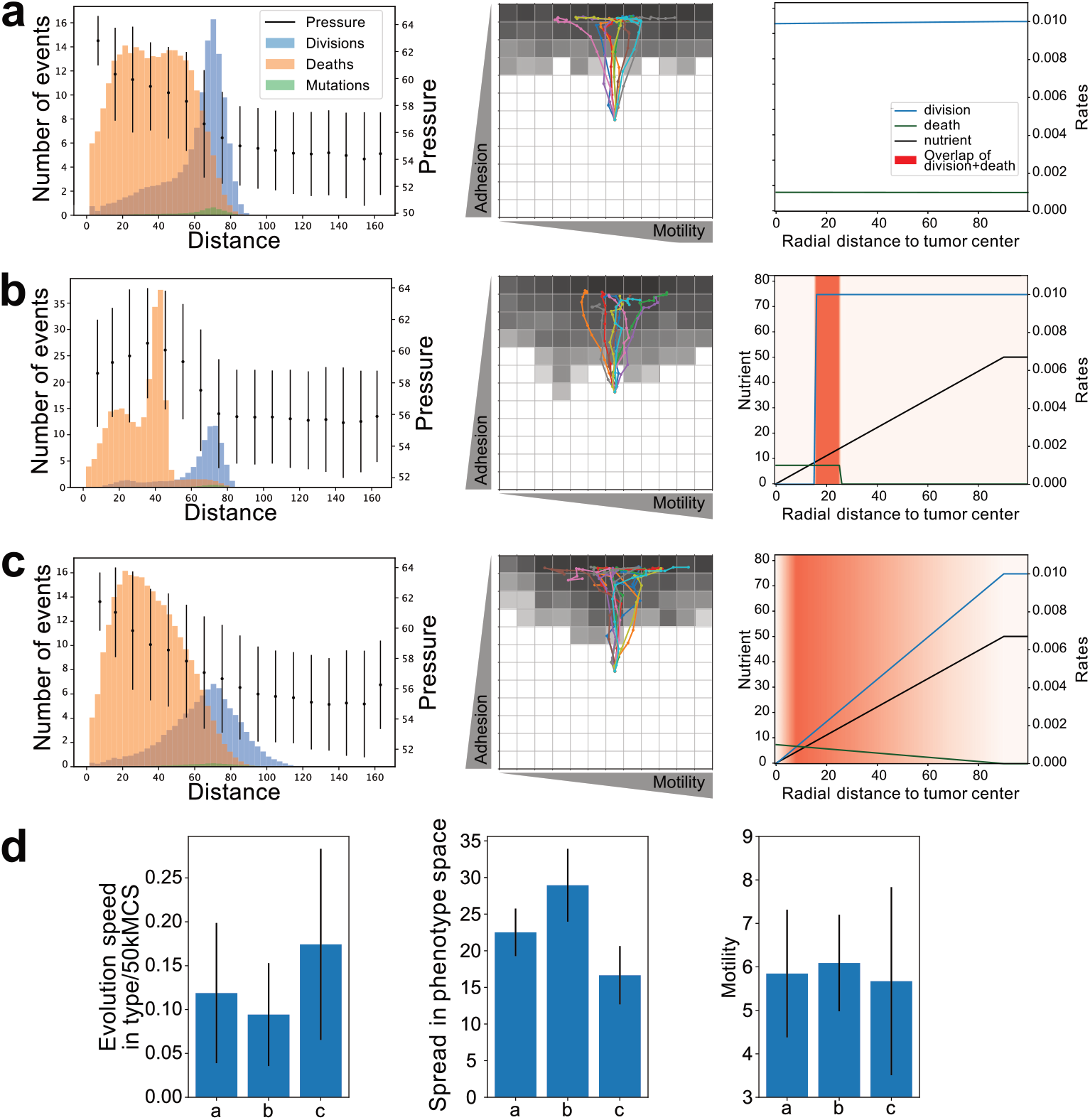
Spheroid tumor growth for different nutrient dependency mechanisms. In a-c, on the left, the radially averaged location of the events with respect to the center of the spheroid on the left. The number of events is normalized with 1*/r*^2^, the non-normalized plots can be found in the SI. Simulation size is 200 voxel^3^. The center plots show the trajectories of the centroid of the phase space occupation. The points on trajectories indicate temporally equidistant points. The shading shows the average distribution in the phenotype space at the endpoint at *t* = 1500 kMCS of 15 simulations. The right plots depict the spatial dependency of the division and death rates for a cell and the dependence on nutrient availability, depending on the distance from the tumor center. The red shading indicates areas of overlap of dividing and dying cells, here evolutionary pressure is most effective. **a** without dependency of cell division and death on nutrient surrounding, **b** division and death regulated by thresholds on the constant nutrient surroundings and **c** linear dependence of the rates on the nutrient. **d** Macroscopic tumor properties for the respective nutrient availability cases of **a**, **b** and **c**. The black lines describe the standard deviation and the bars the mean values of 15 simulations.

Observing the statistical occurrence of cell events in relation to the radial distance to the tumor center, we find that cell deaths are located in the tumor center while cell divisions are located at the tumor margin (see Fig. 2 a). This leads to an inward movement of cells (as seen by the stripes in Fig. 1). There is a buildup of pressure inside the tumor, this is facilitated by the inwards movement of the cells and the central potential. This behavior is well studied in experiments and computational models [28].

The simulation is started with a single cell type in the center of the phenotype space. Through mutations during tumor growth, more cell types are introduced which leads to a distribution in the phenotype space around the initial cell type. With the progressing simulation, the distribution moves in the phase space towards an evolutionary favorable position. The center of mass, spread, and movement speed of this distribution are analyzed and compared. With the model introduced so far, we find that the tumor evolves towards the low adhesion regime. The individual trajectories start with a high directionality towards low adhesion, the trajectories then develop without directional preference along the motility axis (see Fig. 2 a-c).

#### Nutrient Dependency

We investigate how nutrient dependency of cell division and death influences the tumor properties in our model. By introducing a dependency of cell division and death on nutrient availability, cells compete over space and nutrients. Nutrients are introduced as a growth-limiting factor (see Methods), representing, e.g., oxygen or glucose. In an *in vivo* tumor, nutrients diffuse into the tissue and are internally degraded and used up. Here, we simplify his dynamic nutrient surrounding and it is implemented by introducing nutrient availability solely dependent on the position of the cell. The nutrient availability of cells is linearly increasing from the tumor center up to a maximal value and stays constant for further distances, as pictured in Fig. 1 b. We introduce two different dependencies of the cell division rate and death rate on the nutrient availability, as seen in Fig. 2 a-c, right (see also Methods). Firstly, a threshold-based dependency (TBD) that introduces a constant division probability if a cell exceeds a certain nutrient value, and a step decrease in death rate above another threshold. Secondly, we introduce a linear rate dependency (LRD). Here, cells linearly adapt the division and death rates depending on the local nutrient concentration.

Macroscopically, the introduction of a nutrient dependency affects the evolution speed of the tumor (see Fig. 2 d). TBD decreases the evolutionary speed of the system while LRD accelerates the evolution. Enhanced directionality is visible in the different sizes of the spread in the phenotype space. A smaller spread is achieved through a more pronounced directionality of evolution. TBD increases the spread while LRD lowers it. This increase of the evolutionary speed of LRD over TBD can be explained by the larger regions, in which cell divisions and cell deaths occur. In these critical regions (tagged red in Fig 2 a-c, right), the competition between different cell types is most pronounced since cells that can stay in this area or escape outside will survive, while cells that are pushed to the inside will die. Despite the different evolutionary speeds and spreads, the evolution is highly directional towards low adhesion for both mechanisms and no nutrient dependency. The composition center of mass in the phenotype space is only insignificantly changed by the introduction of nutrient dependencies of cell death and division on the migration axis. Interestingly, the introduction of nutrient dependency in a constant environment does not introduce significant evolutionary queues towards higher motility or higher division rate for the parameters chosen here. The nutrient dependency of cell death and division leads to a shift of cell deaths towards the inside of the tumor. TBD introduces a drop in pressure at the center of the tumor (see Fig. 2 b).

The mechanism of nutrient dependency with LRD shows a larger selective pressure on the tumor and therefore leads to a higher speed of evolution. Furthermore, a linear dependency is biologically more reasonable, since cells do not binarily up- or down-regulate cell division in most cases, but adapt continuously [29]. Hence, LRD is used in the subsequent manuscript if not explicitly stated otherwise.

#### Gradient steepness

Next, we observe how the gradient steepness which is linear to the gradient extent influences evolution. Evolutionary speed increases with decreasing steepness of the nutrient gradient(see Fig. 3 a, left). The speed is increased because the extent of the critical area, in which cells both die and divide expands together with the size of the gradient. Therefore, a shallower gradient leads to a larger volume with high selective pressure, which accelerates the evolution. The spread of the phenotype ensemble decreases with shallower gradients. With small nutrient dips, the number of cells inside the gradient is smaller, leading to a larger portion of cells that are outside the gradient. Cells outside the gradient find optimal conditions for growth and therefore do not experience evolutionary pressure. This leads to a loss of directionality of evolution and therefore to an expansion in all directions. A shallower gradient introduces a higher variability in the evolution of the motility/division axis. The median value is not significantly changed. For all subsequent simulations, a *r*_dip_ = 90 is used, and this simulation is indicated by the dashed line in Fig. 3 and 4.

**Fig 3.**
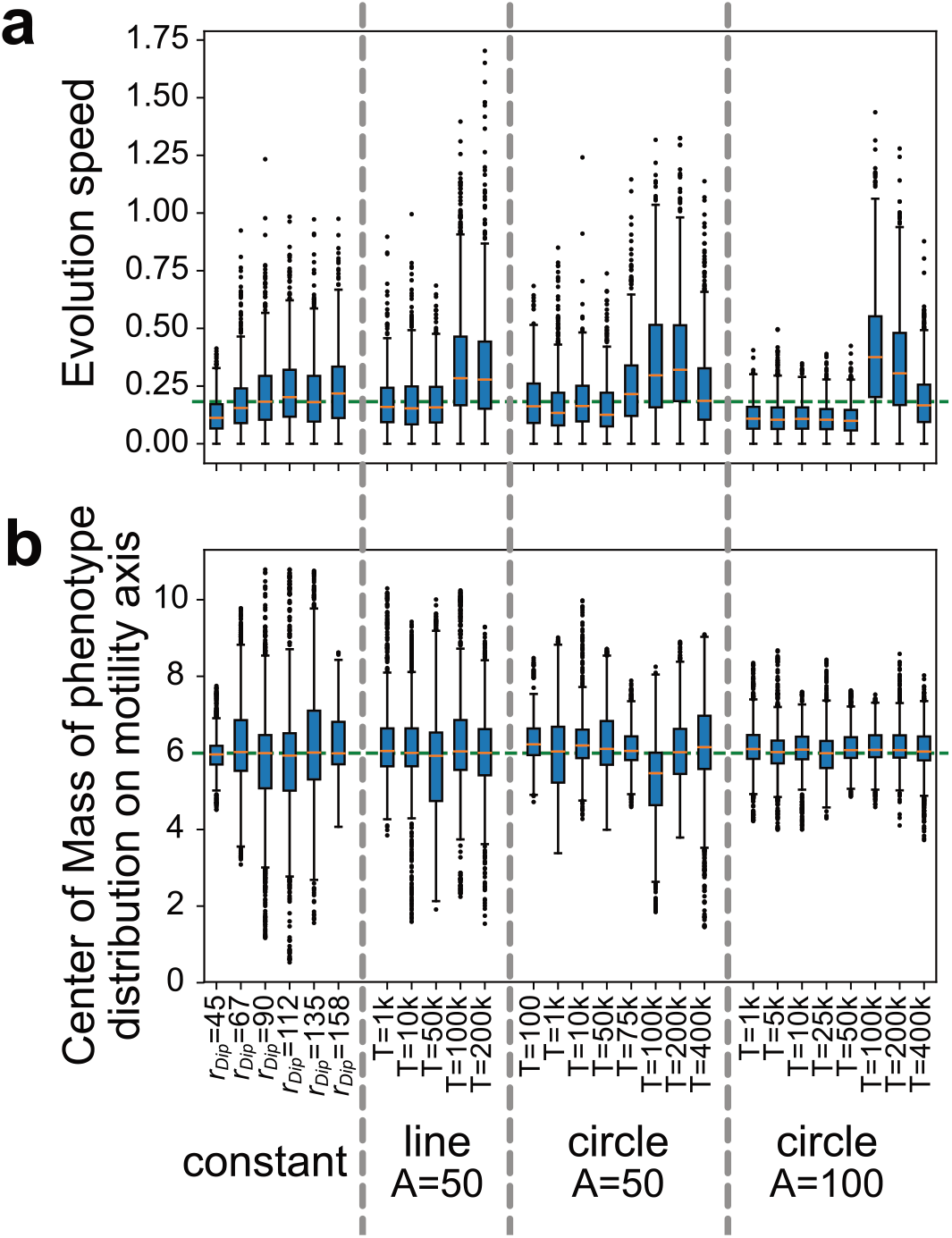
Macroscopic tumor properties for linear dependency of division rates to nutrients. **a** (top) Evolution speed in conformation space. Values are averaged between 1500 − 3000 kMCS. In the first group (left), the simulations are performed with constant nutrient availability and a varying radius of the dip in nutrient availability (*r*_dip_). In the following groups, the nutrient availability is dynamic, and the dip moves on a circle or a line in the x,y-plane with radius *A* and period *T*. In the dynamic cases is *r*_dip_ = 90. The green dashed line indicates the value of the reference simulation with constant surroundings. **b** The average center of mass in the conformation space on the ‘motility–1/division-rate’ axis. Cases are distinguished and data is collected as in a).

**Fig 4.**
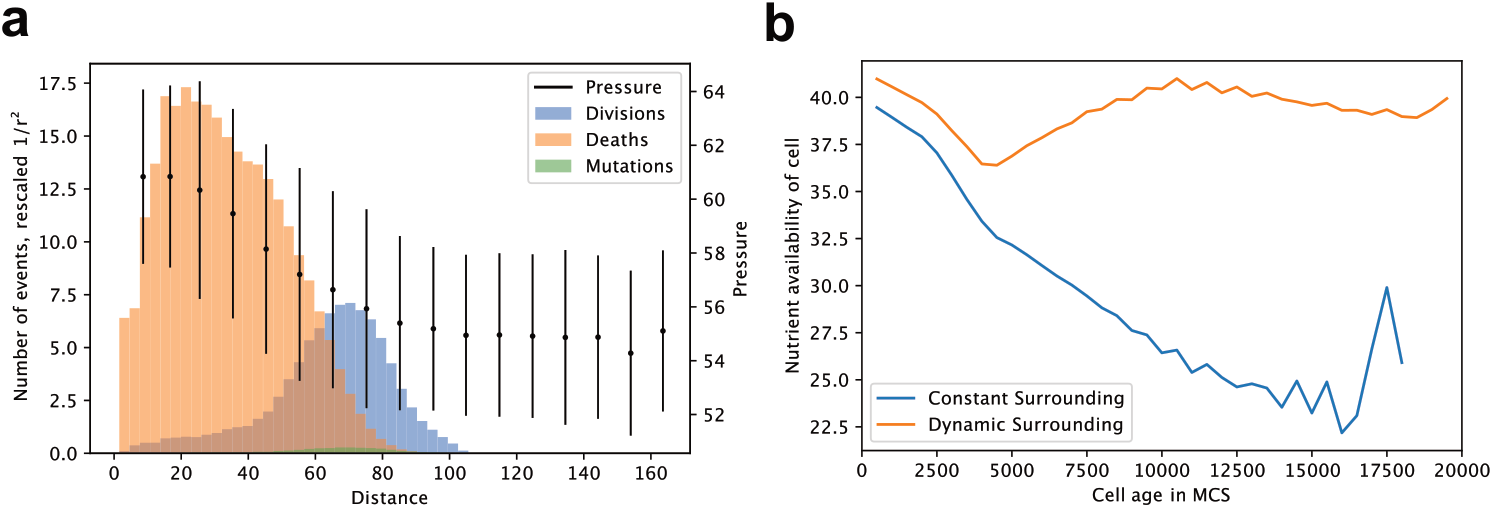
Single-cell properties in dynamic surrounding. **a** Normalized distribution of the locations of cell division, deaths and pressure for dynamic surrounding (line, *A* = 50*, T* = 100 kMCS). **b** Nutrient availability of the cells in relation to the cells age (time after cell division), for constant surrounding and dynamic surrounding (line, *A* = 50*, T* = 100 kMCS).

### Dynamic environment

To examine the effects of a dynamic surrounding, a temporal dependency on nutrient availability was introduced. The center of the negative source was shifted in a periodical manner in one or two dimensions and different extents. All other parameters of the simulation remain the same. Three different possibilities are compared. One, the center of the dip moves on a straight line along the x-axis, following a sine function with amplitude *A* = 50 and period *T*. Two, the center follows a circle in the xy-plane with an amplitude A and period T, here two different amplitudes *A* = 50 and *A* = 100 are compared.

#### Global tumor effects

##### Evolution speed

For small values of the movement period (1 kMCS ≤ *T* ≤ 50 kMCS) the speed is significantly decreased, compared to the constant surrounding case (see Fig. 3 a). By further increasing the period, an increase in evolution speed can be observed for periods between *T* = 75 kMCS and *T* = 200 kMCS. For further increasing values, the evolution speed remains high and decreases for a high amplitude circle case. This increase of evolution speed can be seen independently of the amplitude and motion type of the dynamics at the same frequencies.

##### Composition center

The period *T* of the dynamics does not seem to significantly affect the location of the evolutionary optimum. For the circular dynamics with a high amplitude (circle, *A* = 100), the variability of the optimum is reduced, visible through a smaller standard deviation of the result. The change on the motility axis when introducing a dynamic surrounding is small and can not be conclusively confirmed by only observing the center of mass in the phenotype space (see Fig. 3 b).

##### Spread

For small amplitudes (*A* = 50) the spread is increased over the constant surrounding case. The changing condition at each position presents temporally changing nutrient availability and therefore different evolutionary optima. This leads to a broadening in the phenotype space.

For simulations in which the amplitude of the circle is larger than the nutrient dip (*A* = 100, *r*_dip_ = 90), the spread is elevated significantly more (see SI Fig. 6 c, right). Here, the spread of the distribution in the phenotype space is doubled, compared to the constant case for small values of the movement period (1 ≤ kMCS *T* ≤ 50 kMCS). For larger values of *T* the spread decreases (cf. SI Fig. 6 c). The tumor center is never affected by the nutrient drop, and therefore always provides optimal conditions. This ‘save spot’ in the tumor center is responsible for the increased spread since the evolutionary direction is lost in there.

#### Local cell effects

In Fig. 4 the dynamics and temporal behavior of individual cells are observed.

Tumor cells statistically move from the spheroid boundary inwards towards the tumor center during their lifetime (cf. SI Fig. 11 c). In the constant case, the cells move inwards quicker and die earlier than in the dynamic case. Comparing the nutrient availability of individual cells over their lifetime, a static decline proportional to the distance is visible for the constant case (cf. Fig. 4 b). This is reasonable since the nutrient availability is directly coupled to the position. Looking at the dynamic case, the nutrient availability first decreases, but then increases. This is due to the fact that cells divide in a region in which other cells die, cell death is introduced by a shortage of nutrients. After division, cells move inward towards the tumor center and the nutrient drop continues to move. Here, preferably cells survive that quickly exit the nutrient drop, or divide at the ‘rising edge’ of the moving gradient. We hypothesise that this is the main driver of the change in evolutionary optimum on the motility scale since fast-moving cells have a statistical advantage in this case.

We define the lifetime of a cell as the time between the last division and cell death. The mean lifetime and the extent of the lifetime distribution, significantly increases with the introduction of a dynamic surrounding that enables a ‘save spot’ (cf. SI Fig.12 a). Here, the nutrient dip moves around the tumor in a circle that is larger than the tumor. The nutrient availability in the tumor center is therefore always optimal, providing zero evolutionary pressure and therefore a ‘natural reserve’ on the population scale. We observe the development of the mean lifetime of the tumor cells over time, see SI. Fig. 6 e. Overall, an increase in the mean lifetime is visible for both dynamic and constant nutrient surroundings. This increase in lifetime can be linked to the adaptation of the cellular properties to the surroundings and is a result of the evolution of the system.

#### Fitness evolution

We measure the fitness of each cell by tracing the lineage for the following eight generations and counting the descendants. The acquired numbers are then projected on the cell types and then on the parameter space spanned by the adhesion and motility parameters. Like this, the fitness optimum can be tracked over time. The behavior on the adhesion axis has been identified clearly, all tumors developed towards the low adhesion regime. The motility parameter was less conclusive by observing the center of mass of the cell type ensemble (cf. Fig. 3).

The fitness of the cell types is averaged along the adhesion axis. The resulting distribution shows the fitness in relation to the motility and division rate. This distribution has a clear maximum that is determined by fitting a Gaussian distribution to the data. The top value of this distribution is then plotted over time for different simulations in Fig. 5 a. We assumed to find a clear trend towards high motility cells in a dynamic surrounding, however, the behavior can not be decisively found and may be lost in fluctuations c.f. Fig. 5 b. A trend is visible when comparing the dynamic case of a circular motion with a radius of 100 to the dynamic case that shifts the fitness optimum towards more motile cells at the cost of a lower division rate.

**Fig 5.**
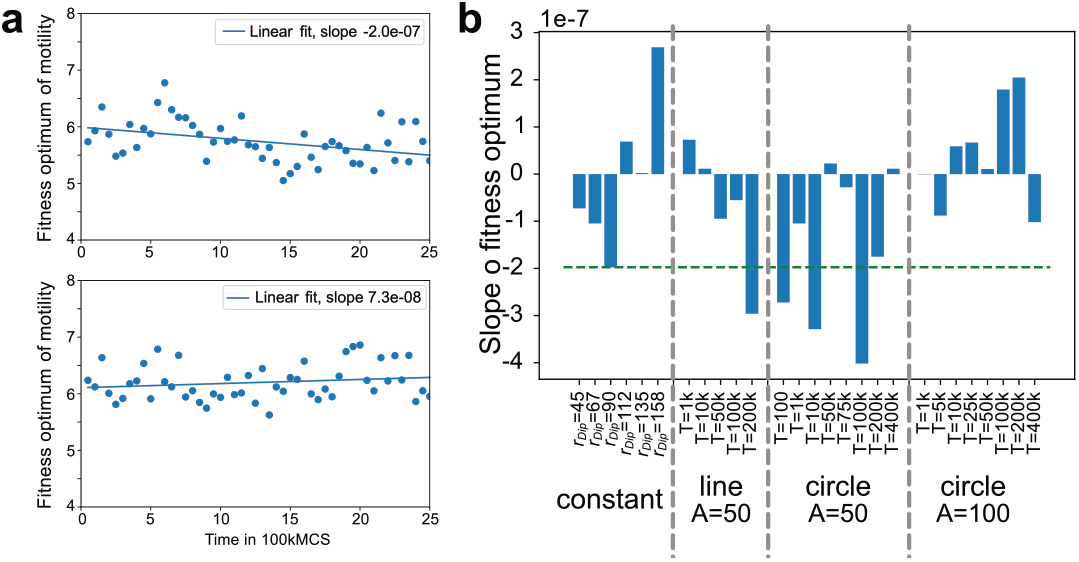
Fitness evolution. The number of descendants in the next 8 generations, is defined as the fitness of a cell. **a** The fitness of all cells is determined at different times and projected on the cell types and their parameters. A Gaussian function is fitted to the resulting distribution and the central value is determined to find the fitness optimum on the parameter axis. Here, the fitness optimum of the motility parameter with division rate trade-off is plotted over time. An average of 15 simulations is pictured. Two simulations are compared, top: constant surrounding, bottom: dynamic on a line, *A* = 50, *T* = 1 kMCS. **b** The slope of the fitness development is plotted for each simulation. A positive slope leads to a development towards high motility, whereas a negative slope leads to a development towards low motility and high division rates.

**Fig 6.**
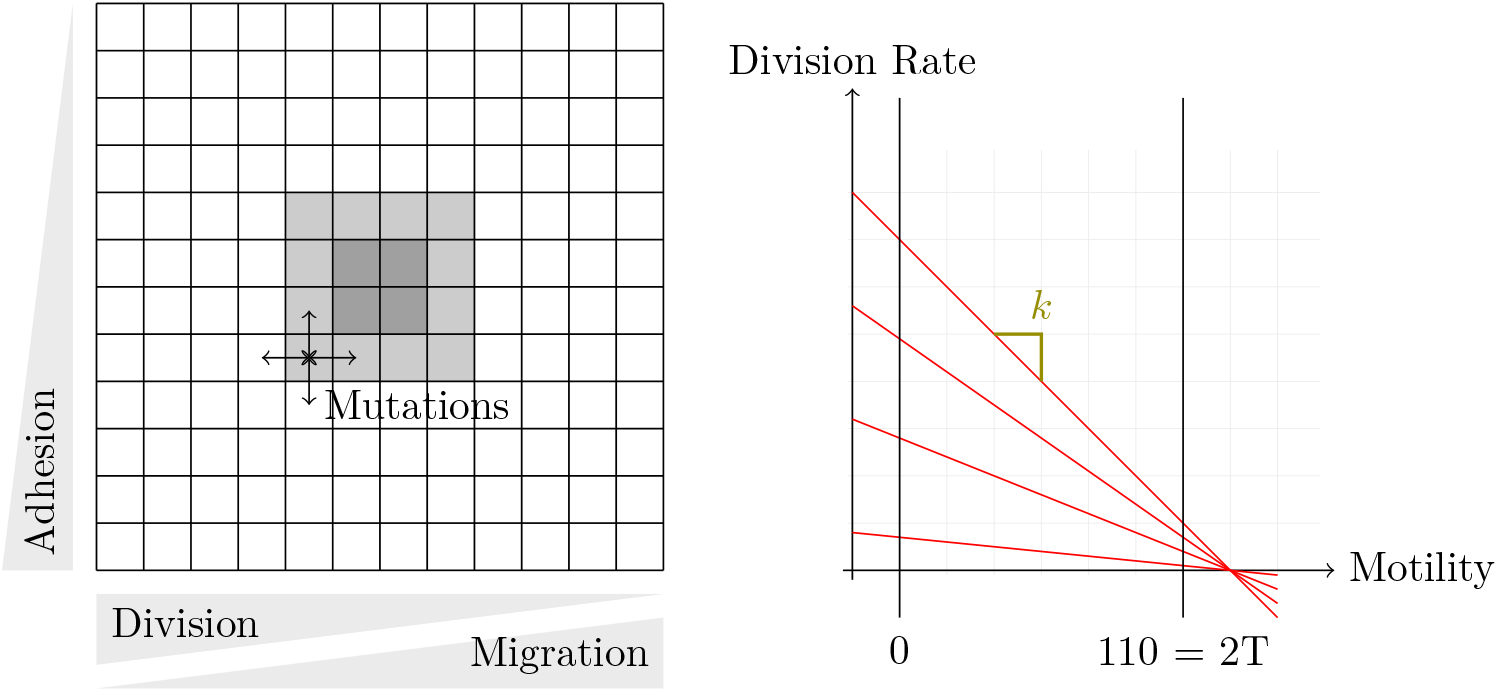
Cell type parametrization in a 12 × 12-matrix. Linear trade-off model between division rate and motility.

## Conclusions

We present a computational model of a spheroid tumor in surrounding tissue. Mutation of cells is enabled by a change of phenotype at cell divisions. Two parameters can be changed during a mutation, cell–cell adhesion and the motility of the cell. We introduce a division rate trade-off for motility. The system is allowed to evolve freely and the tumor composition is tracked in parameter space over time.

We find that the mechanical and geometrical properties of the system are sufficient to drive the ensemble towards low-adhesion cell types. This mechanical effect in tissue evolution has been described in [16]. Mechanical properties alone drive proliferation at the tumor edge and cell death in the center.

We introduce a dependency of cell divisions and deaths on nutrient availability, which is linearly decreasing towards the tumor center. Using a linear dependence on nutrient availability for proliferation and inverse linear dependence for cell death leads to a higher evolutionary speed than a threshold-based dependency.

In *in vivo* tumors, solid stresses through tissue displacement are built up that are able to compress and block blood vessels [22]. This can lead to fluctuating and nutrient availability in tumors. We investigate how fluctuating nutrient availability influences tumor evolution.

We find that a temporally variable nutrient surrounding introduces a larger life span for cells. Especially a ‘save spot’ enables much longer lifetimes and a broader evolutionary spread

We find a significant dependency of dynamic nutrient surroundings on the evolutionary speed in phenotype space. The speed shows a frequency dependency, with a lower evolutionary speed for fast fluctuations followed by an increase over the constant case for lower frequencies. A critical time scale exists for the fluctuation of nutrient availability that provides a distinct peak in evolution speed, which we find to be between *T* = 100 kMCS and *T* = 200 kMCS.

The fitness of cells can be determined by lineage tracking and the fitness is linked to the cell types and their parameters. With this, the effect of parameter values of a cell on its fitness can be determined. While a clear preference in fitness is visible for low adhesion cells, no clear change in the preferential direction of evolution can be identified along the motility axis. A trend towards higher motility is visible for large radii of nutrient fluctuations. We predicted that dynamic nutrient availability influences the fitness optimum of tumor evolution, this could not be conclusively be answered and has to be explored further by extending the range of possible motility and introducing different trade-offs.

Experimental work on spheroid tumors could provide a verification of the results found here. The single-cell motility of cells grown in different spheroid cultures could be measured and compared [30]. The nutrient surrounding of the growing spheroid culture can be varied from a constant availability to a periodically changing nutrient concentration in the surrounding media. We expect to find cells with higher motility in the latter setup.

## Materials and methods

**Table 1.** Parameters varied during simulations.

| Parameter | Range | Dependency |
| --- | --- | --- |
| Cell-cell adhesion | $[0 \dots 2T]$ (no repulsion) | Independent |
| Motility | $[0 \dots 2T]$ | Trade-off motility vs. division rate |
| Division rate<br>$k \in [1e-4 \dots 1e-6]$ | $[l]R = -k(\text{Motility} - 130)$ | |
| Metropolis Temperature $T$ | 55 (Constant) | |
| Motility recalculation Time $t_{\text{motility}}$ | 100 MCS (50,200,400) | |
| System size | $50,100,400 \mu m^3 \approx \text{voxels}^3$ | |
| Coupling to central potential | -70 |  |

Energy functions are:

- Volume
- Surface
- Adhesion
- Random motility
- Central potential

### Adhesion

The cell–cell adhesion is proportional to the contact area between cells and independent of the duration of the adhesion. The strength is not limited or quantized by focal adhesion but only determined by the adhesion parameter between the cell types and the shared area.

### Nutrient

The nutrient availability of a cell is determined by its location in 3D space. The position of a cell is defined as the center of mass of its spatial extent. The function is a radially linear decay within a sphere, in the center of the simulation box. The center of the nutrient well can be temporally constant or moving, to represent constant or dynamic tumor environments. The nutrient represents a growth-limiting factor for the cells.

### Central Potential

To avoid all cells accumulating in the outer regions with constant high nutrient availability, a potential is introduced. This potential leads to all tumor cells experiencing a constant force towards the center point of the simulation. This point is also the center of the nutrient well, with the lowest availability. The potential leads to an increase in pressure at the center of the tumor.

### Random motility

Motility is implemented by assigning a preferential direction of movement to each cell. This direction is defined by a potential along a vector. The three-dimensional direction of this vector is randomly reassigned in a regular interval of 100 Monte Carlo sweeps. The cells are coupled to this potential by a constant force that is determined by the coupling of the energy term to the potential. This coupling constant varies for different cell types and is referred to as the motility strength in this manuscript.

### Cell division and death

To divide, cells need to exceed the age of 2 kMCS and their volumes have to exceed 90% of their goal volume. Similarly, cells can divide once they exceed the age of 4 kMCS.

There are three different cases for dependency of cell division and death on nutrient availability:

1. No dependence: Constant division probability of division () and death ()
2. Thresholds:

- Division: Constant rate (≈0.005) if nutrient exceeds 15, otherwise no divisions
- Death: If nutrient is below 25 higher rate (0.01) otherwise lower rate (0.00001)
3. Linear dependence:

- Division: Rate linearly increases from 0 to ≈ 0.005 with increasing nutrient availability
- Death: Rate linearly decreases from 0.001 to 0 with increasing nutrient availability

Division rates vary from cell-type to cell-type since they are determined by the division rate - motility trade-off.

The tissue that surrounds the tumor does not participate in cell death or division. The cells that make up the surroundings, therefore participate in the entire simulation and act as a medium that the tumor cells at the tumor edge can interact with and redistribute forces and pressure.

### Evolution speed and spread

The speed of evolution is calculated by tracking the center of mass of the distribution of cell types in the phenotype space. The spread is measured by the extent of the distribution.

## Acknowledgments

The authors gratefully acknowledge the Gauss Centre for Supercomputing e.V. (www.gauss-centre.eu) for funding this project by providing computing time through the John von Neumann Institute for Computing (NIC) on the GCS Supercomputer JUWELS at Jülich Supercomputing Centre (JSC).

